## Supplementary Material for "The transformation of sensory to perceptual braille letter representations in the visually deprived brain"

**Supplementary Tables**

Suppl. Table 1. Participant information……………………………………………………………..3

**Supplementary Figures**

Suppl. Figure 1. fMRI searchlight results for within hand braille letter classification. ……………………………………………………………………………...……….………4

Suppl. Figure 2. EEG results for decoding of stimulated hand in time ……………………..…………………………………………………...…………………5

Suppl. Figure 3. EEG time decoding results when only using occipital and parieto-occipital electrodes ……….……………………...…………………………………………...…..6

Suppl. Figure 4. EEG-behavior RSA decoding results when only using occipital and parieto-occipital electrodes ………………………………………………………………………7

Suppl. Figure 5. EEG time decoding results with empirical baseline ……………………...…………………………………………………………...………8

Suppl. Figure 6. Representational similarity of braille letters in neural and behavioral measures using Jaccard similarity ……………………………………………………………...…9

Suppl. Figure 7. Confusion matrices for ROI decoding within hand…………………………..10

Suppl. Figure 8. Confusion matrices for ROI decoding across hand………………………..…12

Suppl. Figure 9. Confusion matrices for EEG time decoding at indicated time points within hand…………………………………………………………………………………..…14

Suppl. Figure 10. Confusion matrices for EEG time decoding at indicated time points across hand…………………………………………………………………………………...15

| Experiment | ID | Sex | Age | Blindness onset | Etiology | Light perception |
|---|---|---|---|---|---|---|
| fMRI | EB01 | F | 24 | birth | Cataract | yes |
| fMRI | EB02 | M | 35 | birth | Retinopathy of prematurity | yes |
| fMRI | EB03 | M | 36 | 2 years | Retinoblastoma | no |
| fMRI | EB04 | M | 39 | 3 years | Retinoblastoma | no |
| fMRI | EB05 | F | 33 | birth | Retinopathy of prematurity | yes |
| fMRI | EB06 | M | 36 | birth | Unknown hereditary disease | no |
| fMRI | EB07 | F | 53 | birth | Glaucoma | yes |
| fMRI | EB08 | F | 34 | birth | Optic nerve atrophy | yes |
| fMRI | EB09 | F | 30 | birth | Unknown malformation | no |
| fMRI | EB10 | F | 47 | birth | Retinopathy of prematurity | no |
| fMRI, EEG | EB11 | M | 55 | birth | Retinopathy of prematurity | no |
| fMRI, EEG | EB12 | F | 42 | birth | Tapetoretinal degeneration | yes |
| fMRI, EEG | EB13 | M | 55 | birth | Retinitis Pigmentosa | no |
| fMRI, EEG | EB14 | F | 38 | birth | Leber Congenital Amaurosis | yes |
| fMRI, EEG | EB15 | F | 35 | birth | Optic nerve atrophy | yes |
| EEG | EB16 | M | 48 | birth | Retinopathy of prematurity | no |
| EEG | EB17 | F | 61 | 3 years | Cataract | no |
| EEG | EB18 | F | 42 | birth | Retinopathy of prematurity | no |
| EEG | EB19 | F | 29 | birth | Leber Congenital Amaurosis | yes |
| EEG | EB20 | F | 41 | birth | Retinopathy of prematurity | no |
| EEG | EB21 | F | 55 | birth | Retinopathy of prematurity | no |

**Supplementary Table 1**. Participant information.

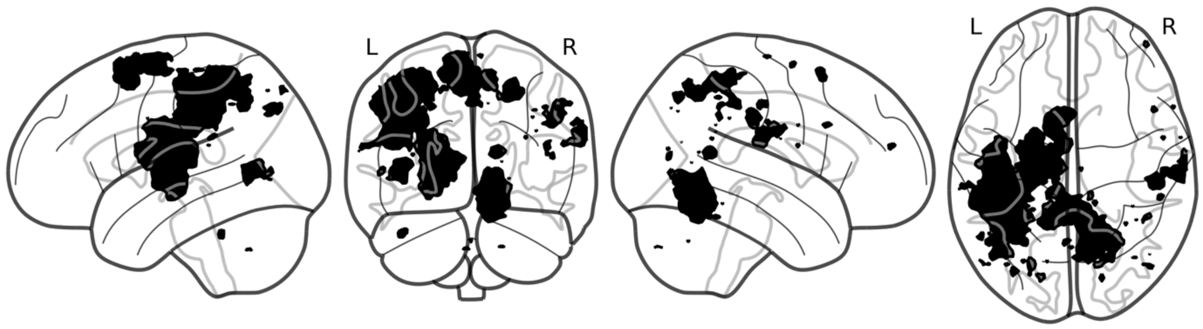


**Supplementary Figure 1.** fMRI searchlight results for within hand braille letter classification. Results index mixed sensory and perceptual braille letter representations (N = 15, height threshold P< 0.001, cluster-level FWE corrected P < 0.05, colored voxels indicate significance).

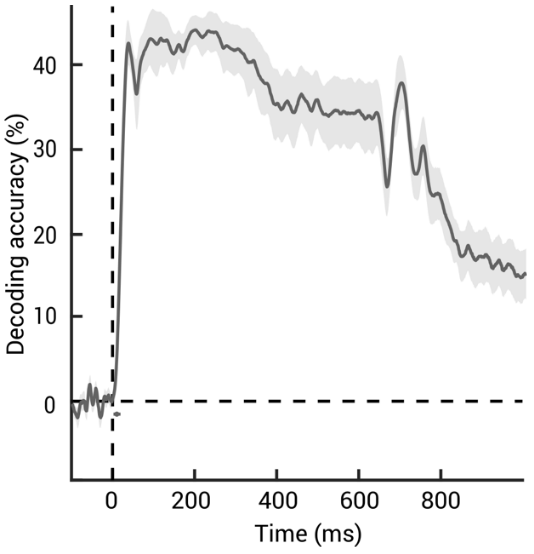


**Supplementary Figure 2.** EEG results for decoding of stimulated hand in time. The shaded area around the curve indicates standard error of the mean. Significance onset and its 95% confidence interval are indicated by a dot and horizontal line below curve (N = 11, 1,000 bootstraps, one-tailed Wilcoxon signed-rank test, $P < 0.05$, FDR corrected).

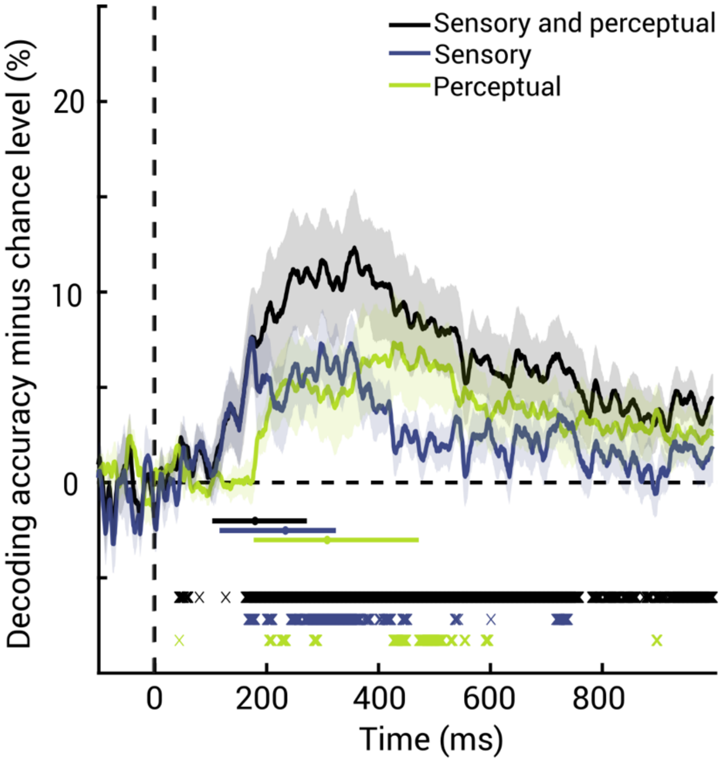


**Supplementary Figure 3.** EEG time decoding results when only using occipital and parieto-occipital electrodes. Analysis is limited to 8 electrodes instead of all electrodes (n=63, see Figure 3a for comparison). For within and across hand decoding, we subtracted the chance level (50%) from the data. The area around curve indicates standard error of the mean. Significance onset and its 95% confidence interval are indicated by a dot and horizontal line below curve (N = 11, 1,000 bootstraps). Significant time points are indicated by x below curves (N = 11, one-tailed Wilcoxon signed-rank test, $P < 0.05$, FDR corrected).

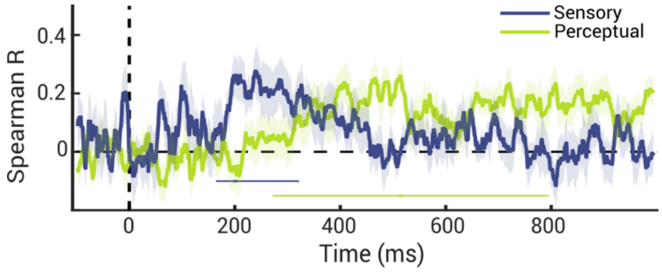


**Supplementary Figure 4.** EEG-behavior RSA decoding results when only using occipital and parieto-occipital electrodes. Analysis is limited to 8 electrodes instead of all electrodes (n=63, see Figure 4c for comparison). Shaded areas around the curves indicate standard error of the mean. Significance onset and its 95% confidence interval are indicated by a dot and horizontal line below curve (N = 11, 1,000 bootstraps, one-tailed Wilcoxon signed-rank test, $P < 0.05$, FDR corrected).

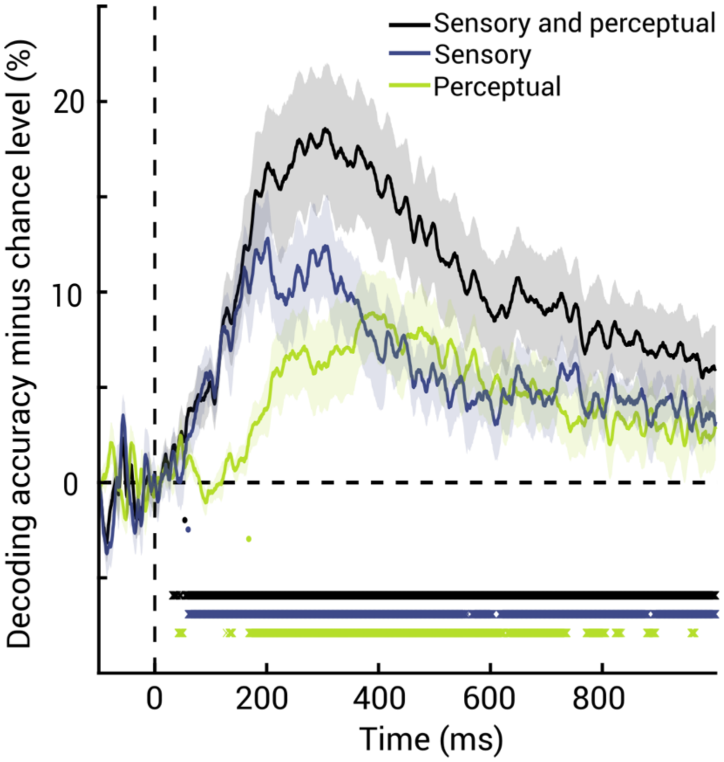


**Supplementary Figure 5.** EEG time decoding results with empirical baseline (see Figure 3a for comparison). For within and across hand decoding, we subtracted the chance level (50%) from the data. Shaded areas around the curves indicate standard error of the mean. Significance onset and its 95% confidence interval are indicated by a dot and horizontal line below curve (N = 11, 1,000 bootstraps). Significant time points are indicated by x below curves (N = 11, sign permutation test,10.000 permutations, $P < 0.05$, FDR corrected).

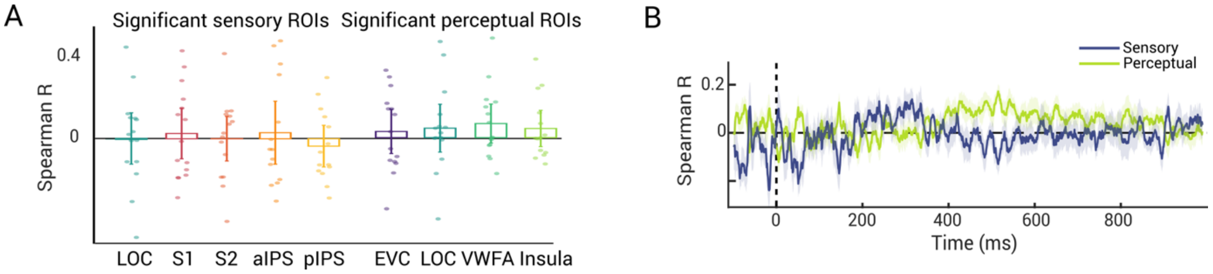


**Supplementary Figure 6.** Representational similarity of braille letters in neural and behavioral measures using Jaccard similarity. A) Results of RSA relating fMRI and behavior in ROIs that showed significant sensory (left) and perceptual (right) braille letter representations in fMRI (see. Fig. 2) (N = 15, two-tailed Wilcoxon signed-rank test, P < 0.05, FDR corrected). Stars below bars indicate significance above chance. Error bars represent 95% confidence intervals. Dots represent single subject data.

B) Results of RSA relating EEG and behavior for sensory (blue) and perceptual (green) representations. Shaded areas around the curves indicate standard error of the mean. Significance onsets and their 95% confidence intervals are indicated by dots and lines below curves (N = 11, 1,000 bootstraps, one-tailed Wilcoxon signed-rank test, P < 0.05, FDR corrected, color-coded as result curves).

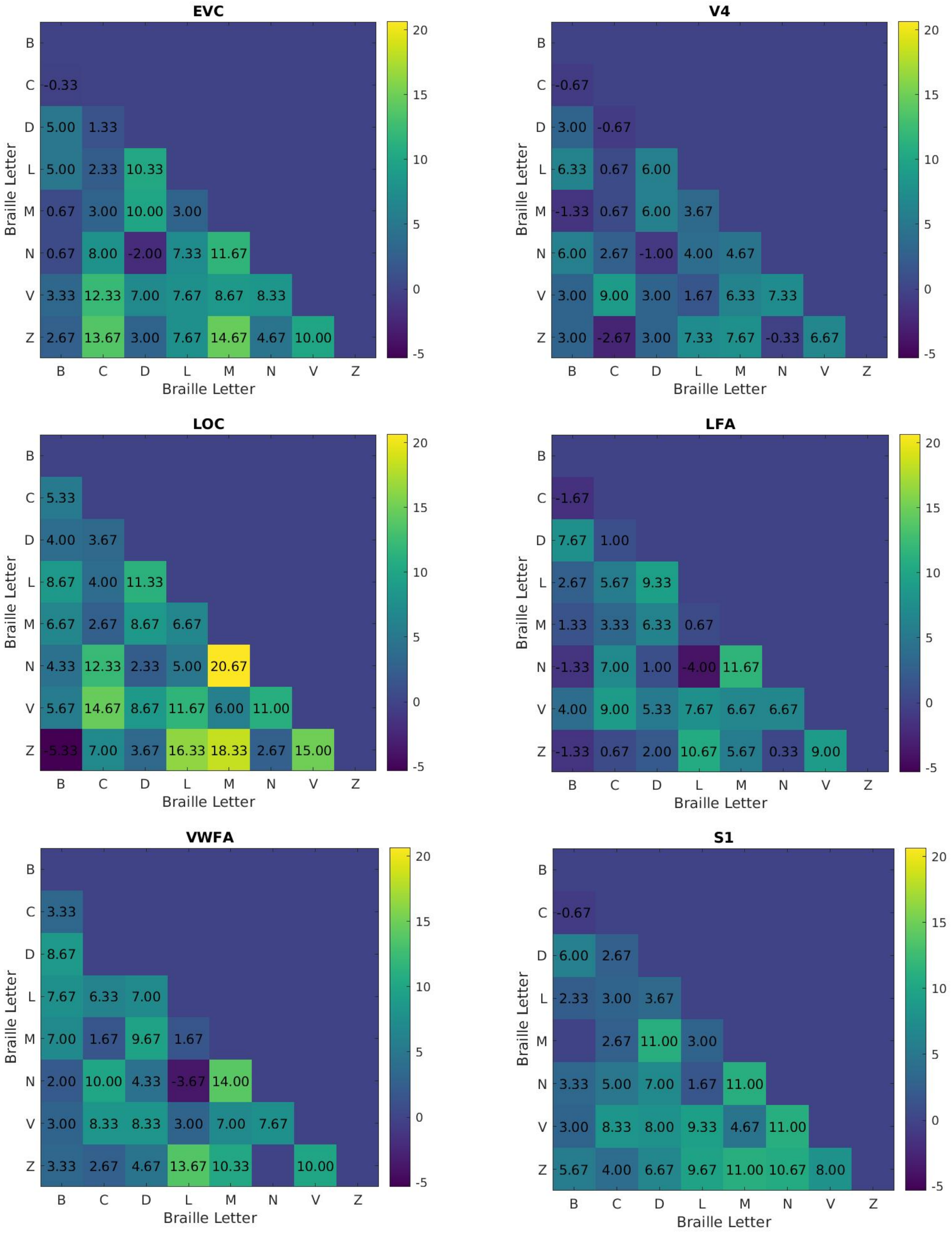
EVC
V4
LOC
LFA
VWFA
S1
Braille Letter
Braille Letter

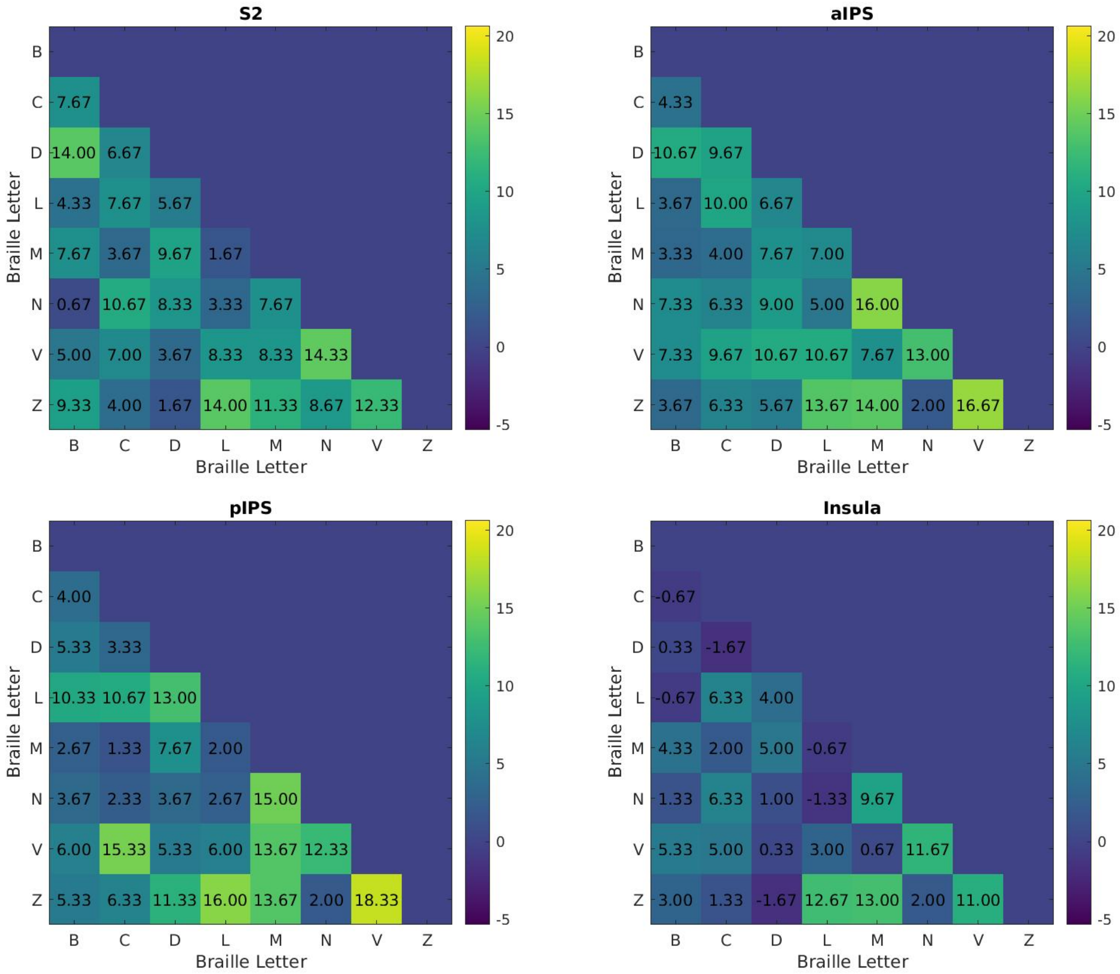


**Supplementary Figure 7.** Confusion matrices for ROI decoding within hand. Values represent pairwise decoding accuracies minus chance level.

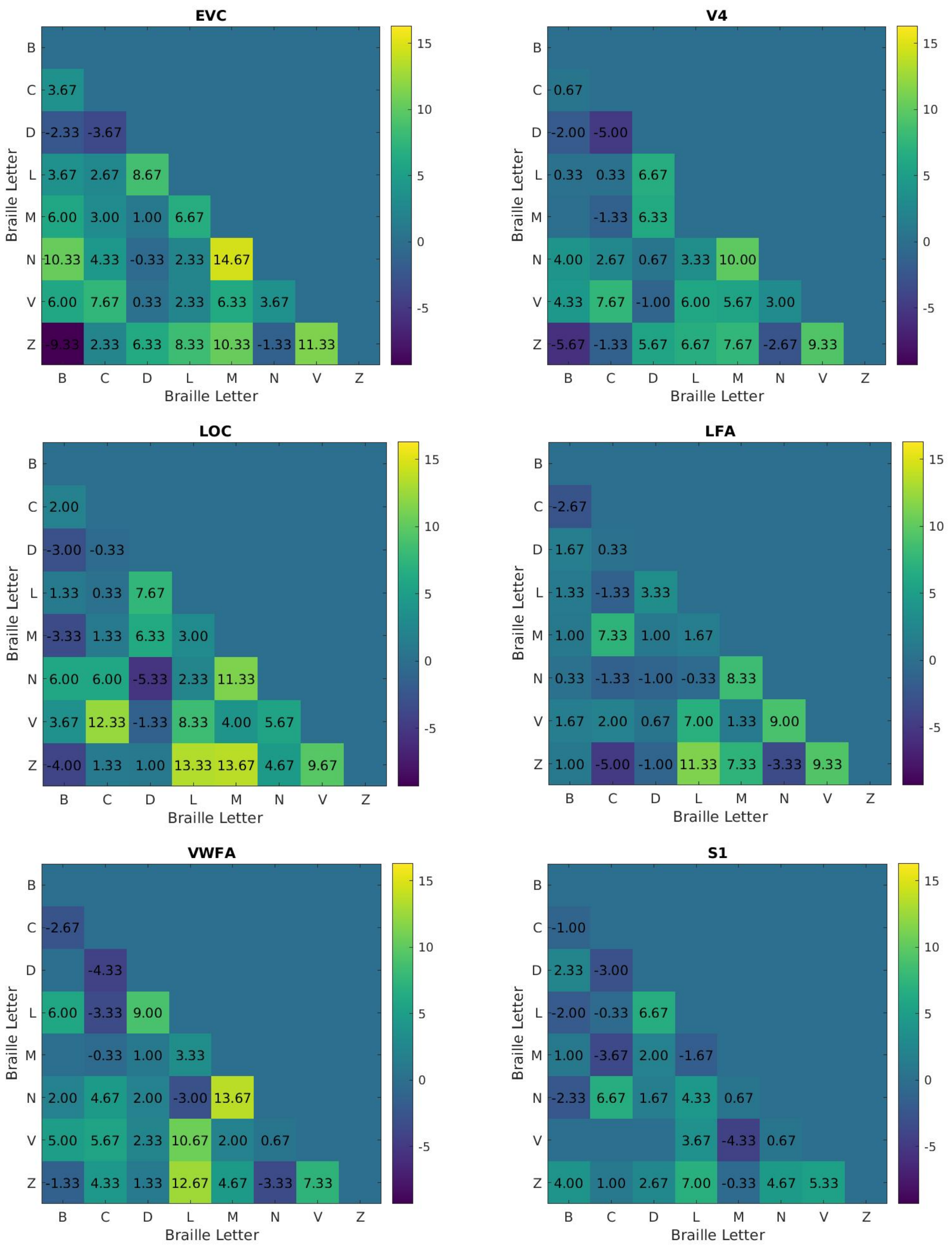
EVC
V4
LOC
LFA
VWFA
S1
Braille Letter
B C D L M N V Z

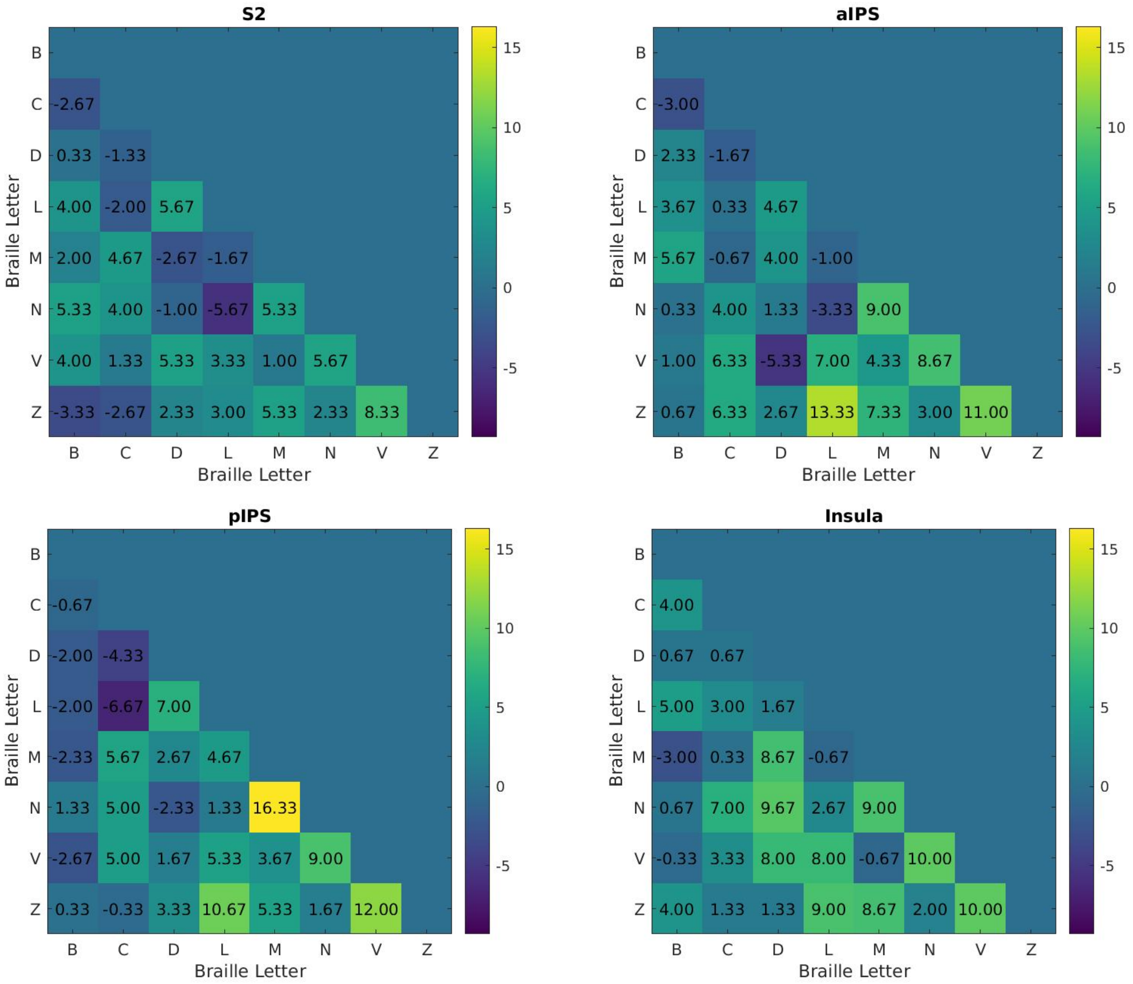


**Supplementary Figure 8.** Confusion matrices for ROI decoding across hand. Values represent pairwise decoding accuracies minus chance level.

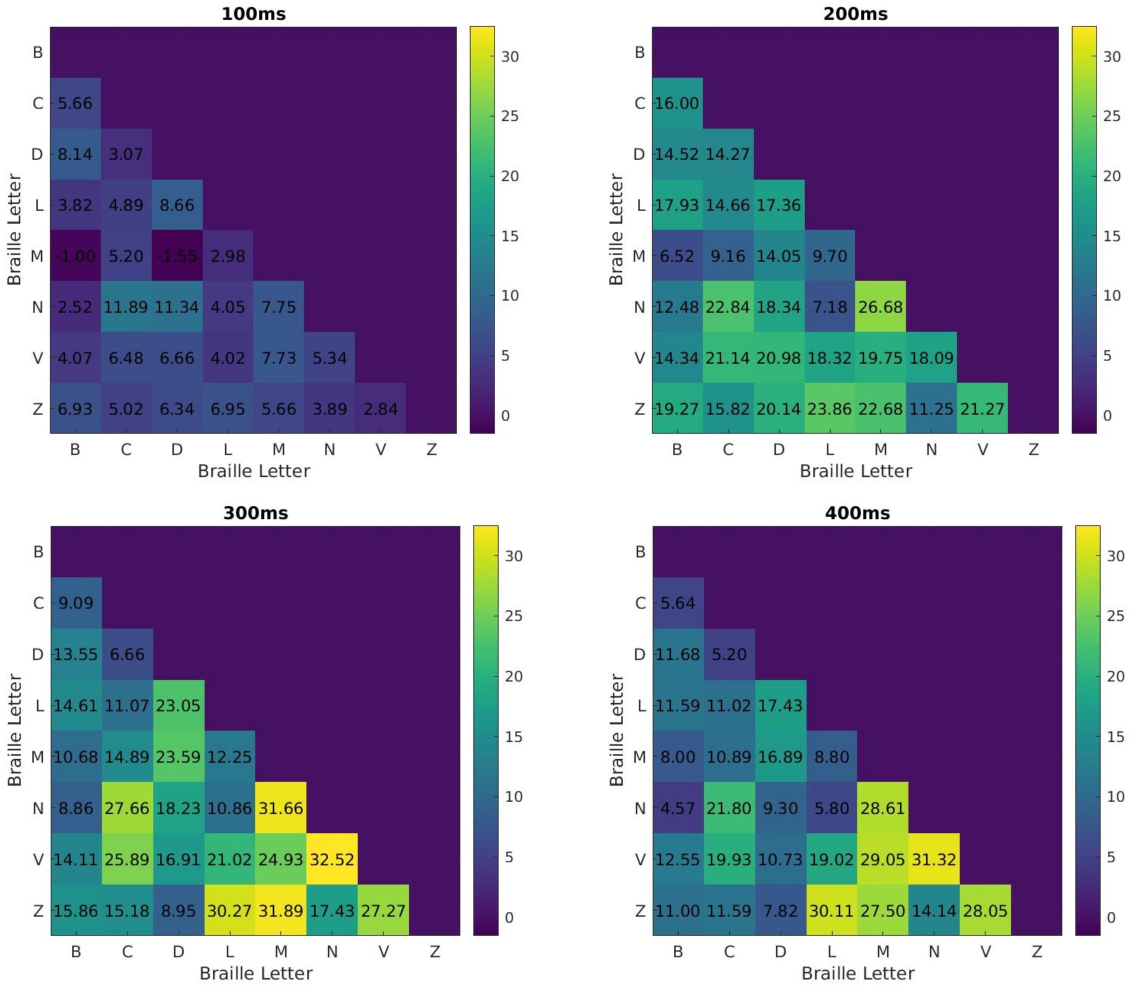


**Supplementary Figure 9.** Confusion matrices for EEG time decoding at indicated time points within hand. Values represent pairwise decoding accuracies minus chance level.

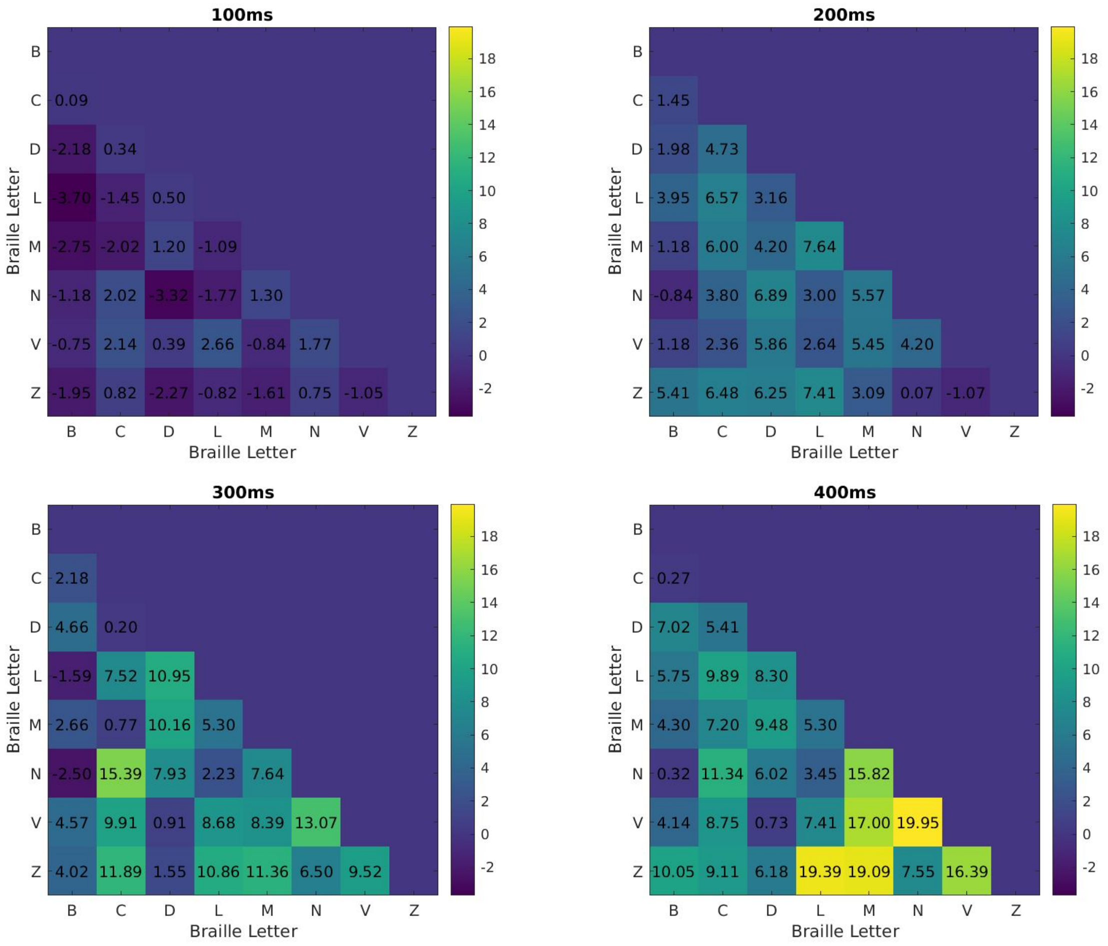


**Supplementary Figure 10.** Confusion matrices for EEG time decoding at indicated time points across hand. Values represent pairwise decoding accuracies minus chance level.
